# Ser68 phosphoregulation is essential for CENP-A deposition, centromere function and viability in mice

**DOI:** 10.1101/2021.11.23.469796

**Authors:** Yuting Liu, Kehui Wang, Li Huang, Jicheng Zhao, Xinpeng Chen, Qiang Wu, Zhouliang Yu, Guohong Li

## Abstract

Centromere identity is defined by nucleosomes containing CENP-A, a histone H3 variant. The deposition of CENP-A at centromeres is tightly regulated in a cell-cycle-dependent manner. We previously reported that the spatiotemporal control of centromeric CENP-A incorporation is mediated by the phosphorylation of CENP-A Ser68. However, a recent report argued that Ser68 phosphoregulation is dispensable for accurate CENP-A loading. Here, we report that the substitution of Ser68 of endogenous CENP-A with either Gln68 or Glu68 severely impairs CENP-A deposition and cell viability. We also find that mice harboring the corresponding mutations are lethal. Together, these results indicate that the dynamic phosphorylation of Ser68 ensures cell-cycle-dependent CENP-A deposition and cell viability.

## INTRODUCTION

Centromere is the specialized chromosomal locus that mediates kinetochore assembly and accurate transmission of replicated chromosomes. In most eukaryotes, centromere identity is not determined by the underlying DNA sequence but the presence of CENP-A-containing nucleosomes (Fukagawa and Earnshaw, 2014).

Interestingly, while the expression and incorporation of canonical histones into chromosomes are coupled to DNA synthesis (Marzluff et al., 2008), the cellular expression and chromosomal assembly of CENP-A is tightly regulated in a DNA replication-independent manner – the mRNA and protein levels of CENP-A peak at late G2 and mitotic phases, respectively (Shelby et al., 2000; Shelby et al., 1997), and the deposition of newly-synthesized CENP-A at centromeresrs at late telophase and early G1 phase of each cell cycle (Jansen et al., 2007; Schuh et al., 2007). In human cells, CENP-A deposition is mediated by its dedicated assembly factor, HJURP (Dunleavy et al, 2009; Foltz et al, 2009).

CENP-A undergoes a variety of posttranslational modifications (PTMs) including phosphorylation, acetylation, methylation, and ubiquitylation (Srivastava and Foltz, 2018). Recent reports have uncovered the importance of these CENP-A modifications in regulating CENP-A deposition at centromeres, the organization of CENP-A chromatin, and the recruitment of CCAN (constitutive centromere-associated network) proteins (De Rop et al., 2012). Among these PTMs on CENP-A, we have previously demonstrated that the dynamic phosphorylation of Ser68 orchestrates the spatiotemporal deposition of CENP-A at centromeres by negatively regulating HJURP recognition (Yu et al., 2015). Our results have also indicated that Ser68 phosphoregulation is crucial for CENP-A stability (Yu et al, 2015; Wang et al, 2021).

In contrast to our findings, another report argued that CENP-A^S68Q^ mutant showed no defects in centromeric CENP-A incorporation and cell viability (Fachinetti et al., 2017). In this report, the CENP-A alleles the authors constructed in the study are intronless, which may compromise the cell cycle-dependent expression and PTMs of CENP-A. Besides, the significance of CENP-A Ser68 phosphoregulation was investigated solely by examining the S68Q mutant CENP-A, which may not fully mimic phospho-Ser68 in the context. Importantly, some of the interpretations of their findings were not accurate, considering that the assembly and maintenance of centromere/kinetochore proteins were still functional in the presence of a small amount of wild-type CENP-A (Fachinetti, 2013, Nature Cell Biology).

To clarify these discrepancies, we constructed CRISPR/Cas9-engineered mammalian cell lines and mice to further assess whether CENP-A Ser68 mutants (S68A, S68E and S68Q) affect centromere functions and cell viability in this report.

## RESULTS

To acutely deplete CENP-A, we generated an R1 cell line with the Myc-tagged osTIR1 stably integrated and expressed. We then introduced HA-AID-tagged wild-type CENP-A into this cell line, allowing the inducible degradation of exogenous wild-type CENP-A proteins upon the addition of auxin (Holland et al., 2012). We further edited the endogenous CENP-A alleles in the cell line to generate point mutations at Ser62, the corresponding residue of human CENP-A Ser68. (Figure 1A). Immunoblot analysis showed that the exogenous HA-AID-tagged wild-type CENP-A was stably expressed along with the endogenous CENP-A (Figure 1B). Upon auxin treatment within 48 hrs, the HA-AID-tagged CENP-A was degraded in cells and no longer detectable by either immunoblot analysis or immunofluorescence microscopy (Figure 1C-D). Furthermore, we verified that the Ser62 coding regions of the endogenous CENP-A loci were successfully substituted to code either Ala62, Gln62 or Glu62 in these R1 cell lines (Figure 1E).

**Figure 1.**
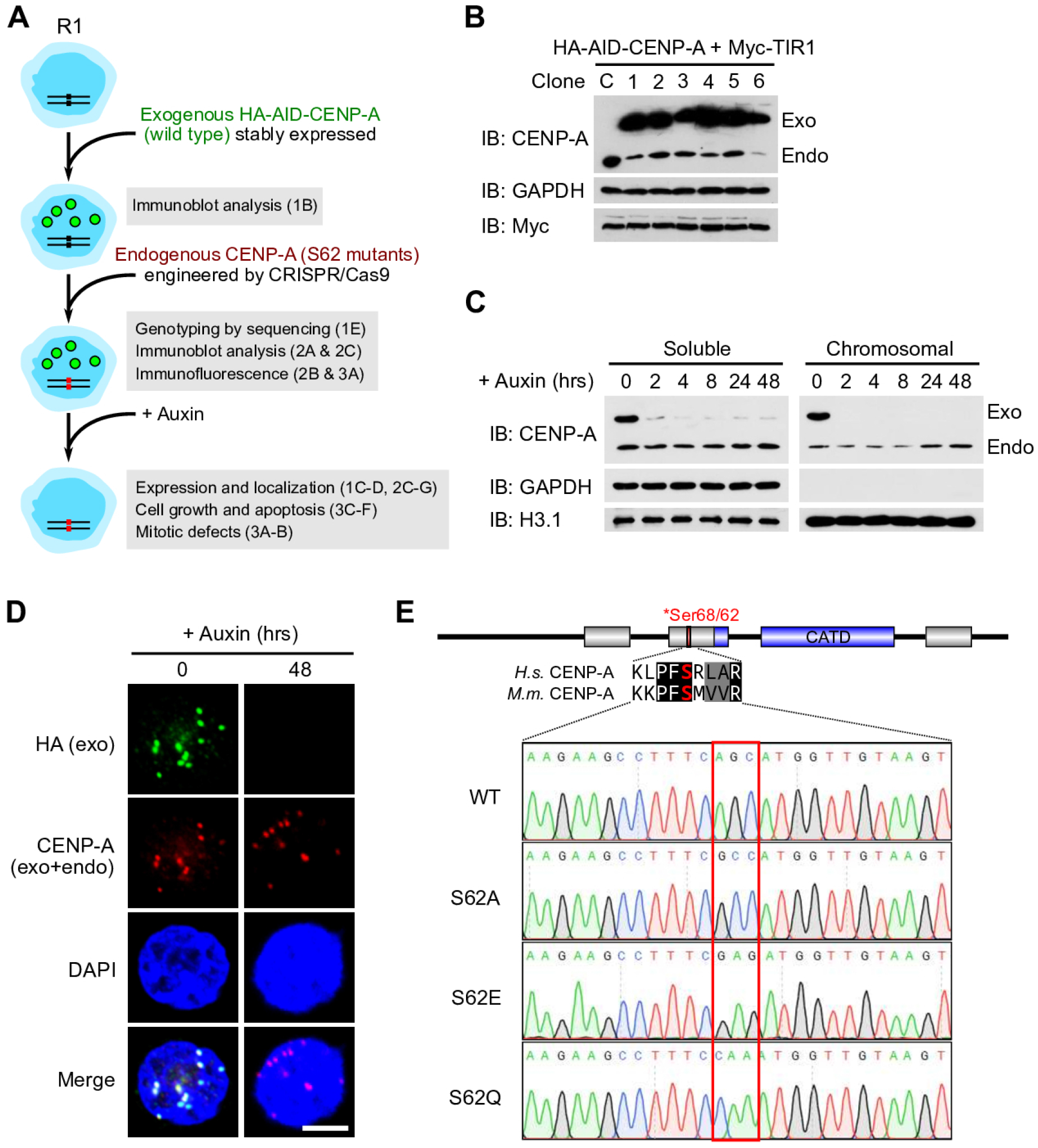
Construction and verification of the CENP-A mutant R1 cell lines used in this study. (A) Schematic of the overall workflow of this study. Results of cell line verification is shown in (B-E). (B) Immunoblot analysis showing six R1 cell lines stably expressing HA-AID-CENP-A and Myc-TIR1 along with endogenous CENP-A. (C) Immunoblot analysis showing that HA-AID-CENP-A protein was degraded within 48 hrs of auxin treatment, from both soluble and chromosomal fractions. (D) Images of R1 cells showing that nuclear HA-AID-CENP-A was degraded upon 48 hrs of auxin treatment. Scale bar, 5 M. (E) Sanger Sequencing results confirm the S62A, S62E and S62Q mutations in engineered R1 cell lines.

Next, we assessed the expression and localization of CENP-A protein in wild-type and Ser62 mutant R1 cell lines. We synchronized cells to G2/M phase and Western blot analysis showed that all cell lines expressed HA-AID-tagged and endogenous CENP-A protein (Figure 2A-B). Importantly, chromatin fractionation assay indicated that only the wild-type and S62A mutant CENP-A could be incorporated into chromatin, whereas the S62E and S62Q mutants mainly existed in the chromatin-free fraction (Figure 2A). By contrast the exogenous HA-AID-tagged CENP-A^WT^ showed enrichment in both soluble and chromatin fractions in all cell lines, and could be readily degraded by adding auxin (Figure 2A-D). Moreover, upon auxin treatment, we also found that endogenous CENP-A^S62E^ and CENP-A^S62Q^ were both unstable and largely degraded from the soluble fraction due to Ser62 phosphorylation-dependent CENP-A degradation (Figure 2C) (Wang, et al, 2021). These results suggested that the mutation from Serine to Glutamate (S62E) or Glutamine (S62Q) impaired the ability of CENP-A loading into centromere chromatin.

**Figure 2.**
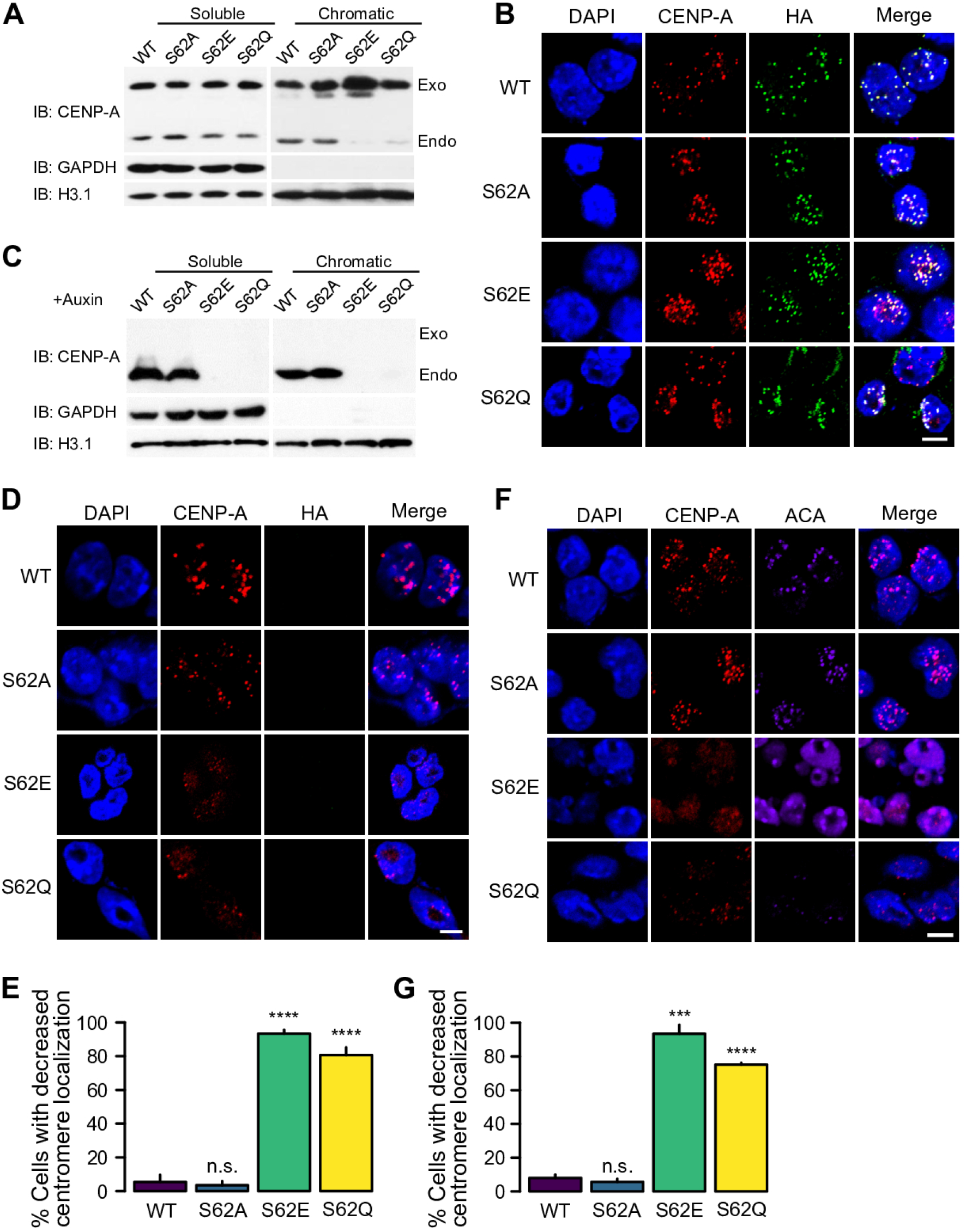
S62E and S62Q mutations impair the centromeric targeting of CENP-A in cultured R1 cells. (A) Western blot analysis showing the expression and distribution of the endogenous CENP-A Ser62 mutants (lower bands) and the exogenous wild-type CENP-A (upper bands) in engineered R1 cell lines. Cells were synchronized to mitosis by adding nocodazole, and were treated with MG132 for 6 hrs before harvest. (B) Images of R1 cells expressing Ser62 mutant CENP-A by immunofluorescence. Scale bar, 5 M. (C) Western blot analysis showing the expression of endogenous S62 mutant CENP-A after 48 hrs of auxin treatment. (D)(F) immunofluorescence images showing the localization of endogenous CENP-A Ser62 mutants after 48 hrs of auxin treatment. HA immunofluorescence indicates the degradation of exogenous CENP-A in (D). ACA immunofluorescence marks the centromeres in (F). Scale bar, 5 M. (E)(G) Quantification of cells with decreased centromere localization in (D) and (F), respectively. n = 149 (WT), 141 (S62A), 152 (S62E), 161 (S62Q) in (E) and n = 150 (WT), 161 (S62A), 170 (S62E) and 165 (S62Q) in (G).

Upon auxin treatment for 48 hrs, we found that the HA-AID-CENP-A^WT^ was degraded in all cell lines and was no longer detected by either immunoblot analysis or Immunofluorescence microscopy (Figure 2C-D). We found that in the absence of wild-type CENP-A, only the S62A mutant CENP-A could be properly assembled into centromeres as the wild-type. Consistent with our previous findings, both S62E and S62Q mutant cell lines showed significantly decreased centromere localization of CENP-A and defective kinetochore assembly (assessed by ACA immunofluorescence) (Figure 2D-G). These findings suggested that CENP-A S62E and S62Q mutations impaired CENP-A deposition and played important roles in regulating CENP-A stability.

We then investigated whether the decreased centromere loading of the S62E and S62Q mutant CENP-A impaired cell division. Our immunoflourescence results showed that the S62E and S62Q mutant CENP-A caused severe mitotic defects, among which lagging chromosomes were observed during mitosis in more than 45% mitotic cells in the mutant cells lines (Figure 3A and 3B). We also wondered whether the abrogation of Ser68 phosphorylation affected cell viability. Therefore, we assayed the ability of cell proliferation of CENP-A mutant cell lines upon acute depletion of the HA-AID-tagged CENP-A^WT^. The cells expressing WT or S62A mutant CENP-A were viable, showing no significant change in the percentage of living cells within 48 hrs of auxin treatment. However, the cells expressing S62E or S62Q mutant CENP-A showed significant cell death starting from 24 hrs of auxin treatment (Figure 3C and 3D). Moreover, we performed flow cytometry (FACS) analysis using annexin V-FITC and PI staining to assess cell apoptosis upon the depletion of wild-type CENP-A in mutant cell lines. The results showed that the substitution of CENP-A Ser62 to either Glu or Gln caused significant cell apoptosis (Figure 3E and 3F).

**Figure 3.**
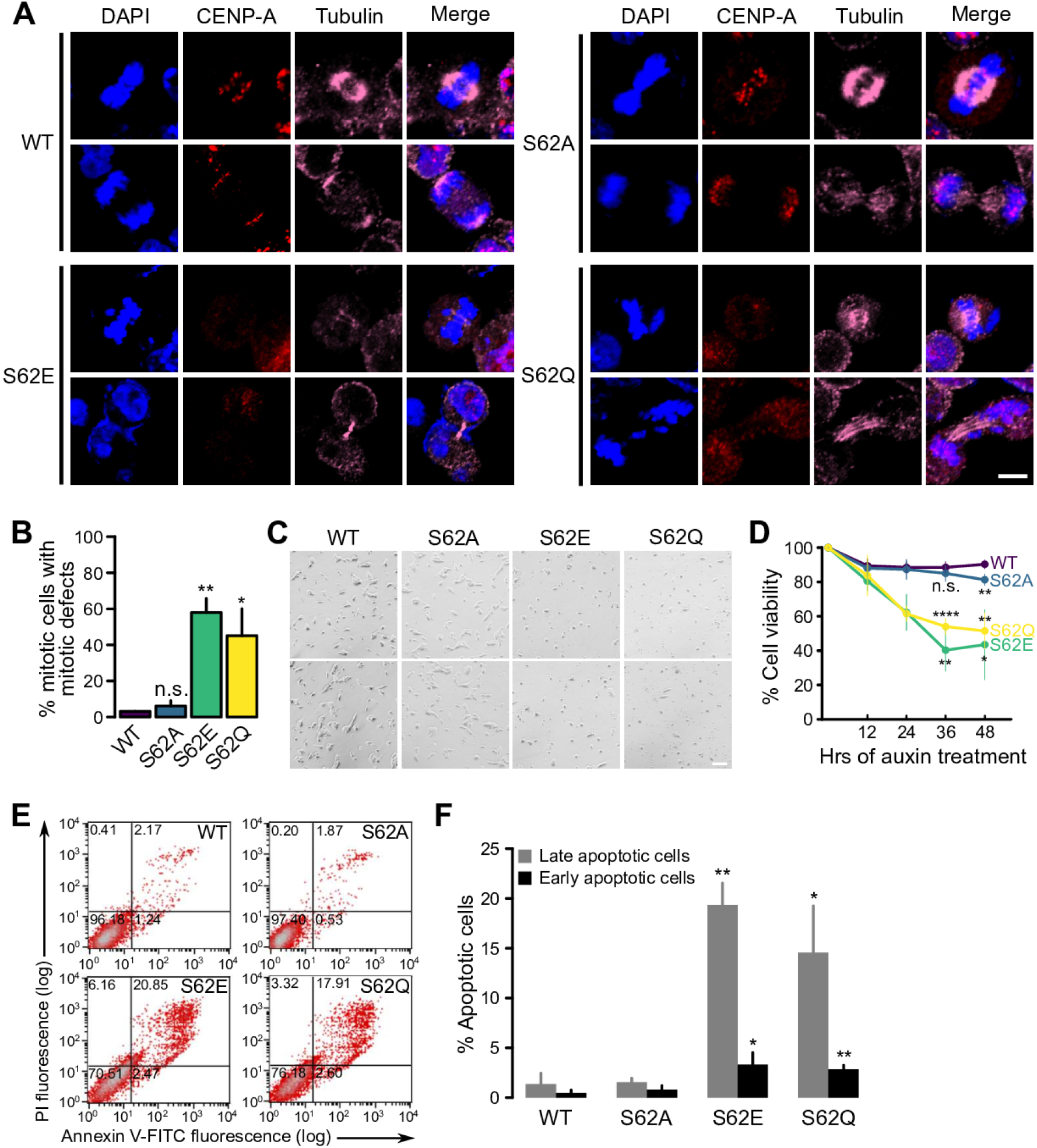
S62E and S62Q mutations lead to increased mitotic defects and apoptosis. (A) Immunofluorescence images showing mitotic defects (lagging chromosomes) of endogenous CENP-A S62 mutant after 48 hrs of auxin treatment. WT or S62 mutant cells were synchronized by adding nocodazole. Scale bar, 5 M. (B) Quantification of (A). The percentage showed the proportion of cells with mitotic defects among all mitotic cells. n = 93 (WT), 98 (S62A), 88(S62E), 89 (S62Q). (C) Microscopic images showing the morphology of WT and S62 mutant R1 cells after 48 hrs of auxin treatment. Scale bar, 100 M. (D) Quantification of the percentage of viable cells in indicated cell lines upon 500 mM auxin treatment. Cells were collected at the indicated time points and stained with trypan blue. Cells were counted on a hemocytometer to calculate the percentage of viable cells based on trypan blue uptake. Mean ± SD is shown for each time point from two independent biological replicates. (E) Apoptosis analysis of WT or S62 mutant cells after 48 hrs of auxin treatment. (F) Quantification of (E). Error bars represent Mean ± SD from two independent biological replicates.

To further validate these observations, we generated CENP-A Ser62 mutant mice using CRISPR/Cas9 (Figure 4A). We identified the genotypes of Ser62 mutant mice by PCR genotyping and Sanger sequencing (Figure 4B). Interestingly, although we successfully obtained multiple homozygous CENP-A S62A mice, no surviving homozygous S62Q or S62D mutant mice were ever obtained despite the fact that we genotyped at least 100 mice for each (Figure 4C and 4E). We thought that the extremely low homozygote rates of S62D and S62Q mice were indicative of a defective embryo viability. Therefore, we further dissected the F2 cross progenies and analyzed the embryos of the S62Q and S62D heterozygous mice at day 10.5 postconception and found empty and dead embryos that were smaller in size compared to wild-type embryos (Figure 4D). Genotypic analysis of the littermates of S62Q and S62D progenies showed no viable homozygotes. Together, these results suggested that the S62D or S62Q mutant mice are embryonic lethal. We also tried to construct CENP-A S62E (which corresponds to CENP-A S68E that also impaired CENP-A deposition in culture human cell lines) mice, but unfortunately, we weren’t able to obtain any mice carrying the S62E mutation after many rounds of microinjections. These results confirmed that CENP-A Ser68 (Ser62 in mice) was indispensable for cell and mice viability.

**Figure 4.**
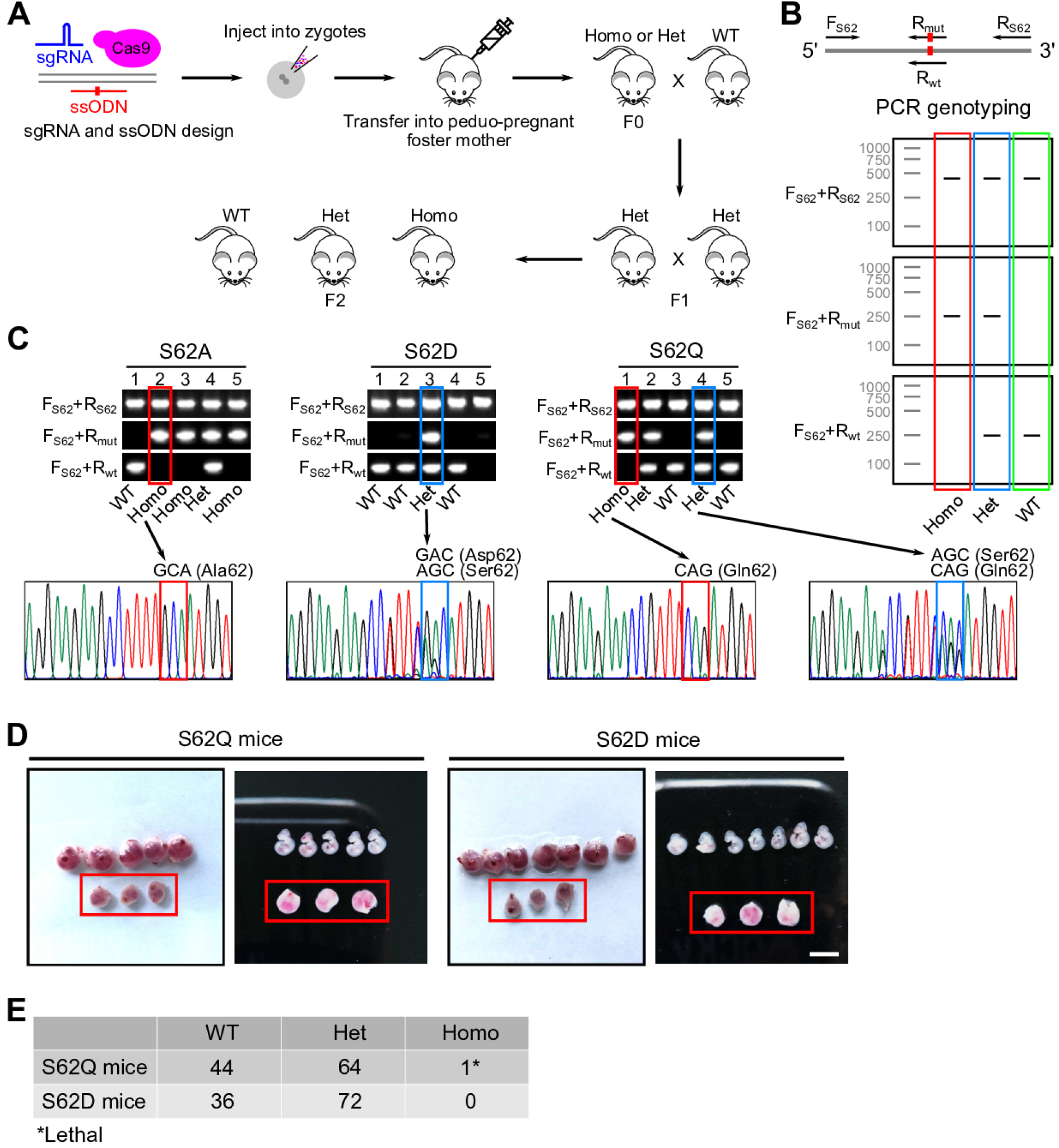
CENP-A S62Q and S62D mutant mice are lethal. (A) Schematic illustrating the construction of mice harboring the CENP-A Ser62 mutations. (B) Primer design and genotyping of CENP-A Ser62-edited mice. Upper, primer pairs that were used for genotyping. The red rectangle indicates the endogenous Ser62 coding region of *Cenp-A*. Lower, anticipated patterns of PCR products for mice of different phenotypes. (C) Genotyping and validation of F2 S62 mutant mice. (D) Embryonic lethality of CENP-A S62Q and S62D mice. Empty embryos at day 10.5 postconception were highlighted in red rectangles. (E) Numbers of genotypes of CENP-A S62Q or S62D mutant embryos from day 9 to day 11.5 postconception.

## DISCUSSION

Our results further demonstrated and extended the conclusions in our previous paper (Yu et al., 2015) - the phosphorylation of Ser68 is necessary for CENP-A deposition at centromeres, and the disruption of the dynamic regulation of pSer68 CENP-A severely affects centromere functions and cell viability.

Fachinetti et al., 2017 have confirmed that S68Q mutant CENP-A was deficient in recruiting HJURP compared to wild-type CENP-A (down to around 25% of wild-type levels; Figure 3B in Fachinetti et al., 2017), and the centromeric staining of the S68Q mutant was significantly weaker and smaller compared to wild-type CENP-A. (Figure 2D, 2F, 2I in Fachinetti et al., 2017). Moreover, a considerable amount of S68Q CENP-A was mistargeted to non-centromeric regions (Figure 1C in Fachinetti et al., 2017). These results are consistent with ours. Our previous report has demonstrated that the phosphomimicking of CENP-A Ser68 impairs CENP-A deposition and results in mitotic defects in cultured human cells (Yu et al., 2015). In line with previous observations, the defective cell viability of Ser62 mutant mouse cell lines and the embryonic lethality of CRISPR-engineered Ser62 mutant (S62D, S62E, S62Q) mice all indicate that the same mechanism (CENP-A Ser68/62 phosphorylation) is regulating CENP-A deposition that is mediated by HJURP in an intact animal model

Different from the system used by Fachinetti et al., 2017, we introduced the Ser68/62 mutations at the endogenous CENP-A loci in cells and animals, which largely maintain the cell cycle-dependent regulation of CENP-A. Together with our previous observations, we demonstrate that the Ser68 phosphorylation ensures accurate cell-cycle-dependent CENP-A deposition at centromeres, and highlight that the dynamic regulation of Ser68 phosphorylation plays key roles in centromere functions and cell viability.

## ACKNOWLEDGMENTS

This work was supported by grants from the National Natural Science Foundation of China (31630041, 31521002, 31991161 to G.L.; 32070604 to J.Z.), the Ministry of Science and Technology of China (2017YFA0504202 to G.L. and 2019YFA0508903 to J.Z.) and the Beijing Municipal Science and Technology Committee (Z201100005320013 to G.L.). The work was also supported by the CAS Key Research Program on Frontier Science (QYZDY-SSW-SMC020 to G.L.), the Chinese Academy of Sciences (CAS) Strategic Priority Research Program (XDB19040202), and HHMI International Research Scholar grant (55008737) to G.L. All fluorescence imaging data were collected and processed at the Center for Bio-imaging, Core Facility for Protein Sciences, Institute of Biophysics, Chinese Academy of Sciences.

## AUTHOR CONTRIBUTIONS

Conceptualization, G.L.; Methodology, K.W., Y.L, L.H., J.Z., and Q.W.; Validation, K.W., Y.L., L.H., and X.C.; Formal Analysis, K.W., Y.L., L.H., Z.Y.; Investigation, K.W., Y.L., L.H., J.Z., X.C. and Z.Y.; Resources, Q.W.; Writing, K.W., Y.L., L.H., Z.Y. and G.L.; Visualization, K.W., Y.L., L.H. and Z.Y.; Supervision, G.L. Funding acquisition, J.Z. and G.L.

## MATERIALS AND METHODS

### Plasmids and DNA sequences

See Table S1 and S2 for plasmids and DNA sequences (oligos, primers) used in this study.

### Cell Culture

Mouse embryo stem cells were cultured in the medium containing 80% DMEM (EmbryoMax, SLM-220-B), 15% FBS (Hyclone, SH30070.03), nonessential amino acids (EmbryoMax, TMS-001-C), 2-mercaptoethanol (EmbryoMax, ES-007-E), l-glutamine (EmbryoMax, TMS-002-C), Nucleosides (EmbryoMax, ES-008-D), Pen/Strep (EmbryoMax, TMS-AB-2C) and 1000 U/ml leukemia inhibitory factor (LIF) (ESGRO, ESG1107) in standard incubator with 5% CO2 at 37°C.

### Western blot analysis

Cells were harvested after 48 hrs of auxin treatment and lysed in RIPA buffer (50 mM Tris-HCl pH 7.5, 350 mM NaCl, 1 mM EDTA, 0.5% Nonidet P-40, 1 mM PMSF, 1 g/mL aprotinin, 1 g/mL pepstatin, 1 g/mL leupeptin). Cell lysates were centrifugated for 15 min at 12,000×g at 4 °C. The supernatant was the soluble fraction. The pellet was washed twice with RIPA buffer and sonicated for 5 mins (3s/8s) after resuspended in buffer containing 50mM Tris-HCl (pH8.0), 10mM EDTA, 1% SDS and protease inhibitors. This slurry was the chromatin fraction. The soluble fraction and the chromatin fraction were denatured by adding SDS sample buffer before analysis of the proteins by SDS-PAGE and Western blot. The primary antibodies used were listed as following: HA (huaxingbio: HX1820) antibody, CENP-A (Cell Signalling Technology, 2048S) antibody, GAPDH (Santa Cruz, sc-25778) antibody, H3.1 (Abcam, ab1791) antibody, Myc (huaxingbio, HX1802) antibody.

### Immunofluorescence and Microscopy

For immunofluorescence assay, cells were grown on glass coverslips, washed twice with PBS and then fixed with 4% paraformaldehyde in PBS for 15 min at room temperature. After three times washes with PBS, cells were permeabilized with 0.1% Triton X-100 in PBS for 15 min. Cells were blocked with 5% BSA in PBS for 1 h at room temperature and then incubated with primary antibodies overnight at 4 °C. Cells were washed for three times with PBS containing 0.1% Triton X-100 and then incubated with fluorescence-conjugated secondary antibodies at 37 °C for 1 h and stained for DNA with 10 mg/ml DAPI (Sigma). The coverslips were sealed with nail polish. Fluorescent images were collected using a 60×oil-immersion lens on an Olympus FV1200 microscope (Tokyo, Japan). For each measurement, at least 50 cells were used to eliminate variations in staining and image acquisition (n=3 experiments).

The primary antibodies used were listed as following: HA (huaxingbio: HX1820) antibody, CENP-A (Cell Signalling Technology:2048S) antibody, ACA (Antibodies Incorporated: Inc.#15-235-0001) antibody,

### Generation of Ser62 mutant R1 cell lines

For generation of mouse ES stable cell line expressing exogenous HA-tagged CENP-A, one day after transient transfection, the cells were seeded into 10-cm dish by serious dilution and were selected by adding Zeocin (Gibco) at concentration of 500 g/mL for one week. Then alone clones were picked and examined by Western blot and fluorescence microscopy. To generate CENP-A Ser62 mutation knock-in cell lines, pX260 was modified to contain the guide sequence insert site of pX330 (Xiong et al., 2018). The donor plasmid containing the homologous arms for recombination was constructed with the corresponding mutations. The homologous arm containing the PAM sequence of SpCas9 target site was mutated to disrupt the PAM sequence. The donor plasmid and the pX260 plasmid were co-transfected into ES cells using Lipofectamine 3000 (Invitrogen) according to the manufacturer’s instructions. Next, cells were seeded into 10-cm dish at low density; and 24 h later puromycin (InvivoGen) was used to select clones for two weeks. Then alone clones were picked out and screened by PCR followed by Sanger sequencing.

### Generation of Ser62 mutant mice

C57BL/6N female mice (4-6 weeks old) were superovulated by injection with 5 IU of pregnant mare serum gonadotropin (PMSG) (Millipore), followed 5 IU of human chorionic gonadotropin (hCG) (Millipore) 48 hrs later. The super-ovulated mice were mated with C57BL/6N male mice and zygotes were collected 20 hrs later from the oviducts. CENP-A Ser62 mutation template DNA was synthesized *in vitro*. To obtain Cas9 mRNA for the microinjection of zygotes, the Cas9 coding sequence was cloned into pcDNA3.1 plasmid under the control of T7 promoter. The plasmid was then linearized by XbaI and used for *in vitro* transcription with mMESSAGE mMACHINE T7 Ultra Kit (Thermo Fisher). After digestion of the DNA template, the transcribed Cas9 mRNA was purified with the MEGA clear Kit (Thermo Fisher). To obtain sgRNAs for microinjection of zygotes, we performed *in vitro* transcription using DNA templates generated by PCR with a forward primer containing a T7 promoter followed by targeting sequences and a common reverse primer. *In vitro* transcription was performed with the MEGA short script Kit (Thermo Fisher) using T7 polymerase by incubating at 37 °C for 5 hrs. The template DNA was removed by digestion with DNase I. The transcribed sgRNAs were purified with the MEGA clear Kit (Thermo Fisher) and eluted in TE buffer (0.2 mM EDTA). Cas9 mRNA (100 ng/ L), Ser62 mutation template DNA (100 ng/ L) and sgRNA (50 ng/ L) each were mixed and injected into zygotes by a Piezo-driven micromanipulator and then cultured in G1 plus medium at 37°C in a humidified incubator with 5% CO_2_ overnight. About 16 hrs later, 2-cell-stage embryos were transferred into the oviducts of pseudo-pregnant ICR female mice. For genotyping, tail tips from mice were lysed in lysis buffer (1% SDS, 5 mM EDTA, 0.4 M NaCl, 20 mM Tris-HCl, 400 g/mL Proteinase K) and genomic DNA was extracted as template for PCR. The sequences of sgRNA and the sequences of the primers used for PCR were listed in Key Resources Table.

### Cell Viability Assay

The Ser62 mutant CENP-A stable cell lines were seeded at 1.5×10^5^cells/well in a 6-well plate, and 500 mM IAA was added every other day. Cells were collected at the indicated time points, stained with trypan blue (Corning), and counted on a hemocytometer to calculate the percentage of viable cells out of the total cells based on trypan blue uptake.

### Cell apoptosis assay

Cells were assayed for apoptosis with the Annexin V-FITC Apoptosis Detection Kit (Beyotime; C1063) and detected by flow cytometry according to the manufacturer’s protocol. All cell samples were analyzed using a FACSCalibur cytometer (BD Biosciences).

### Quantification and statistical analysis

#### Quantification of CENP-A localization at centromeres

The integrated signal intensity of nuclear CENP-A immunostaining were calculated in ImageJ. Centromeric regions were manually segmented and the ratio of intensities of centromeric CENP-A to nuclear CENP-A were calculated. Cells that have the ratio < 0.5 were defined as decreased CENP-A localization. Data from three biological replicates were presented as Mean ± SD.

#### Quantification of mitotic defects

Mitotic cells that were acquired in images from three experiments were collected together to analyze the numbers of mitotic cells that showed defects such as lagging chromosomes, chromosome bridges and multipolar spindles. Data from three biological replicates were presented as Mean ± SD.

#### Quantification of cell viability and apoptosis

Viable cells and apoptotic cells were analyzed as stated in the Methods section. Data from three biological replicates were presented as Mean ± SD.

#### Statistical analysis of the quantification data

In all cases, Ser62 mutant samples were compared to wild type. *p < 0.05, **p < 0.01, ***p < 0.001, ****p < 0.0001 (students’ t-test). n.s., not significant.

---

**Table S1.** Plasmids used in this study.

| Plasmid | Usage |
| --- | --- |
| pcDNA4-HA-AID-CENP-A | Stable expression of exogenous CENP-A |
| pX260-S62 | CRISPR-based Ser62 editing in R1 |
| pblunt-S62A | Donor plasmid |
| pblunt-S62D | Donor plasmid |
| pblunt-S62E | Donor plasmid |
| pblunt-S62Q | Donor plasmid |

**Table S2.**
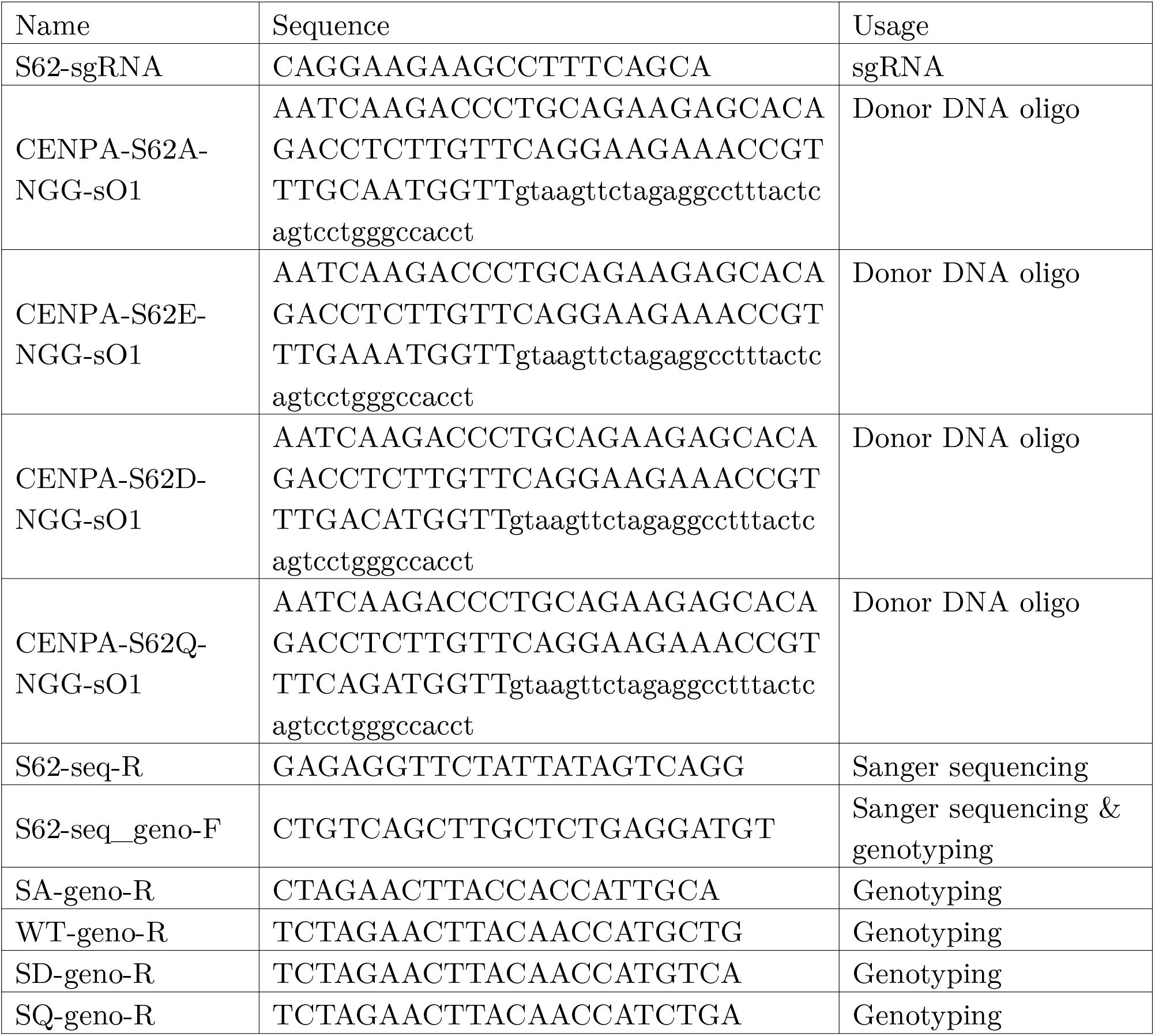
DNA sequences used in this study.

